# Proteomic landscape of multi-layered breast cancer internal tumor heterogeneity

**DOI:** 10.1101/2021.08.05.455361

**Authors:** Mariya Mardamshina, Anjana Shenoy, Daniela Necula, Kateryna Krol, Daniel Pirak, Nitay Itzhacky, Irina Marin, Bruria Shalmon, Roded Sharan, Einav Gal-Yam, Iris Barshack, Tamar Geiger

**Affiliations:** Department of Human Molecular Genetics and Biochemistry, Tel Aviv University, Tel Aviv, Israel; Department of Molecular Cell Biology, Weizmann Institute of Science, Rehovot, Israel; Pathology Institute, Sheba Medical Center, Tel Hashomer, Ramat Gan, Israel; School of Computer Science, Tel Aviv University, Tel Aviv, Israel; Oncology Department, Sheba Medical Center, Tel Hashomer, Ramat Gan, Israel

## Abstract

Despite extensive research, internal tumor heterogeneity presents enormous challenges to achieve complete therapeutic responses. Changes in protein expression are central determinants of cancer phenotypes that reflect potential therapeutic targets. However, previous proteomic studies did not address internal heterogeneity, therefore, masked the necessary spatial resolution to achieve a comprehensive understanding of cancer complexity. Here we present the first large-scale multi-focal breast cancer proteomic study of 330 tumor regions which associated cancer cell function, pathological parameters, and spatial localization of each tumor region. We found marked internal proteomic heterogeneity even within tumors presenting homogeneous receptor expression. Additionally, analysis of the internal heterogeneity, based on coexisting receptor expression or histological patterns in single tumors, showed significant functional differences between homogeneous and heterogeneous tumors related to cancer metabolism, immunogenicity, and proliferation. We anticipate that this study will serve as a starting point towards the development of improved cancer therapy and diagnostics.

## Main

The interplay between cancer genetic aberrations and external microenvironmental cues is the basis of tumorigenic growth, metastasis, and development of highly heterogeneous tumors. Although tumor heterogeneity has been thoroughly investigated at the genomic and transcriptomic levels^1–3^, limited studies have investigated heterogeneity at the proteomic level^4^. Routine breast cancer classification is still based on few molecular markers (i.e., estrogen receptor, progesterone receptor, and HER2); however, limitation to the known breast cancer receptors neglects additional protein determinants of the tumors and even fails to account for the heterogeneous expression of these markers ^5–11^. Here, we aimed to examine the proteomic landscapes within single tumors, irrespective of their clinical subtype, and investigate the importance of the heterogeneous characteristics of the tumors for their progression.

### Spatially-resolved multi-region proteomics

To comprehensively examine proteomic internal breast cancer heterogeneity, and associate cellular functions with histopathological diversity, we performed IHC-guided multi-focal proteomic analysis of 330 histopathologically mapped regions derived from 35 treatment-naive primary breast tumors (Fig. 1A, B; Extended Data Fig. 1A; Supplementary Table 1A-B). Altogether our cohort was comprised of 311 tumor regions, including 12 non-invasive cancer regions and 19 regions from normal adjacent healthy breast tissue, as controls from 13 patients (Extended Data Fig. 1A; nine regions on average per tumor; Supplementary Table 1B). As a preliminary validation of the proteomic data, we examined subtype-specific differences, which recapitulated known characteristics, such as cytokeratin signatures, proliferation, and metabolic functions (Extended Data Fig. 1B-E). Global unsupervised clustering of all regions based on the entire proteome revealed a multilayered complexity of receptor expression patterns, histology, and patient identity. Tumor regions are segregated into four main clusters (Fig. 1C), primarily based on receptor expression (Fig. 1D). Hormone receptor-positive (HR+) clusters were further segregated based on the histological characteristics (Fig. 1E, F). Normal samples were clustered together irrespective of patient identity and closer to the receptor-positive regions, while tumor grade did not have a major effect on clustering (Fig. 1D, E). Interestingly, we found that TN regions that are embedded in HR+ tumors are commonly clustered with their tumor of origin, while several tumors with strong HER2+ expression formed a distinct cluster.

**Fig. 1:**
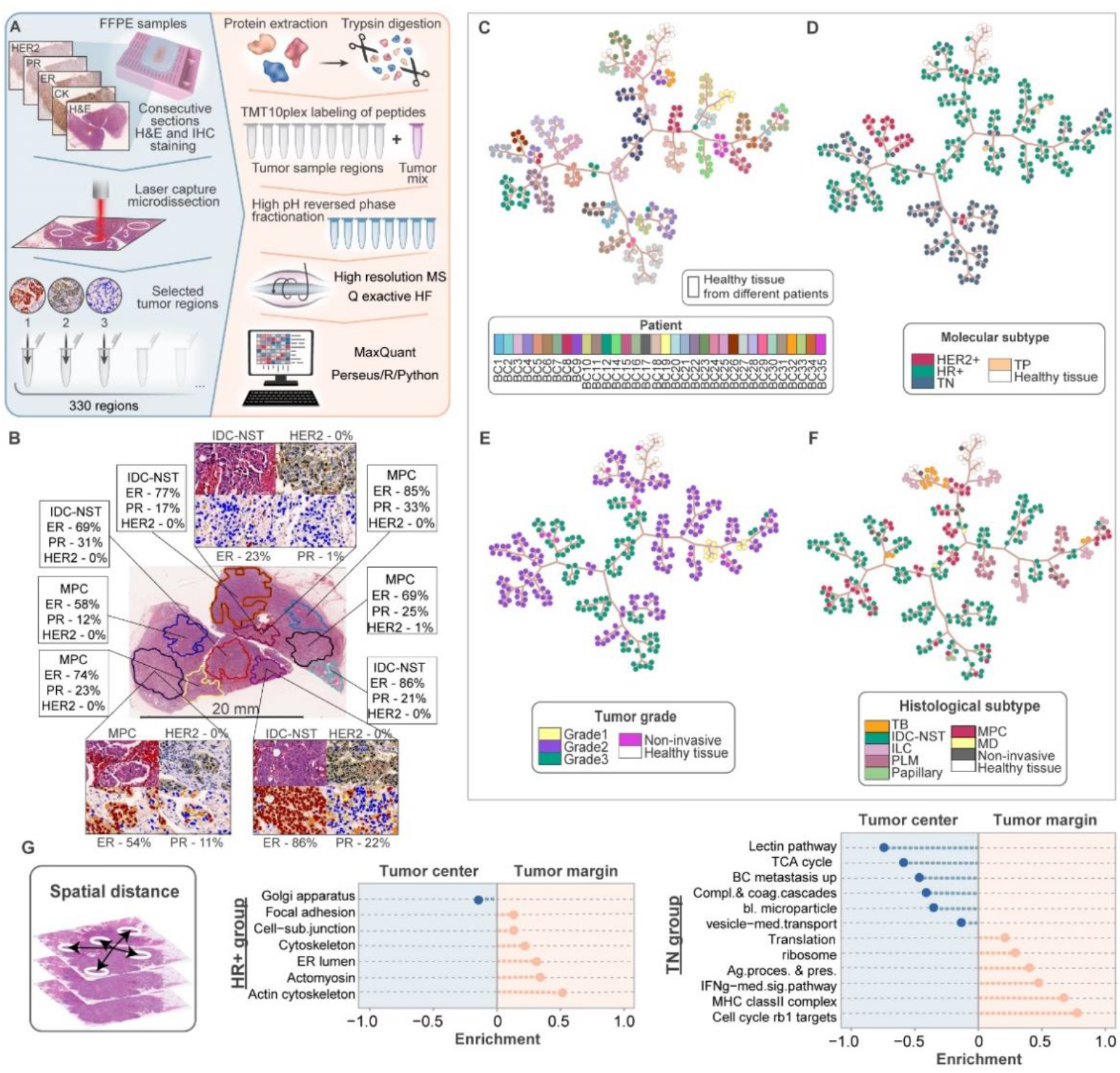
Unsupervised analysis of proteomics of breast cancer. (**A**) Proteomic analysis was performed on FFPE tumor sections. To map the tumor histological and molecular characteristics, we performed immunohistochemistry and H&E staining of consecutive sections and followed with laser-capture microdissection of histopathologically-defined regions from each tumor. Extracted proteins were trypsin-digested, labeled with TMT10plex, mixed with an internal standard, and separated using high pH reverse phase fractionation. Peptide samples were analyzed on the Q Exactive HF MS followed by computational analysis. (**B**) Histopathological analysis of intratumor heterogeneity was based on the combination of ER, PR, HER2, and H&E staining of consecutive sections and quantification using the eSlideManager software. BC7 is shown as an example of the divergence of ER expression and histological features among regions of the same tumor. Images of receptor staining show the overlay of quantification analysis that demonstrate different expression intensity detected by the algorithm, where brown color indicates high intensity staining, orange – moderate intensity, yellow – mild intensity, and blue – no staining. (**C-F**) Tree-and-leaf representation of dendrogram based on modified minimum variance method where each node represents a tumor region, branches of the tree are constructed based on Euclidean distances and the cluster separation is presented by the spatial localization of the branches. Presented dendrograms are color-coded based on: (**C**) Patient identity; (**D**) Molecular subtype; (**E**) Tumor grade; and (**F**) Histological subtype. (**G**) Schematic representation of spatial distances measurement in each tumor. Barplots show enrichment analysis of HR+ and TN tumors. 1D Enrichment analysis was performed with FDR q-value < 0.02.

Supervised analysis of molecular subtype differences showed that the multiregional analysis has a marked advantage over bulk analyses in capturing significantly changing proteins between subtypes, including known subtype markers like *HER2/ErbB2, Muc1, CD44*, and keratins (Extended Data Fig. 2A, B). In addition, by combining the multisampling technique with the physical coordinates of each region within the tumor, we found proteomic determinants of their spatial localization. HR+ regions closer to the tumor margins were enriched for focal adhesion-related proteins like ACTN1 that might suggest higher migratory potential^12^, while peripheral regions from TN tumors appear to be more proliferative with higher antigen presentation than their internal counterparts (Fig. 1G). Furthermore, we found the *FSCN1* and *SRC*, which are known to be associated with cell migration, invasion, and high metastatic potential to be highly expressed in peripheral regions, irrespective of receptor expression^13,14^.

### Quantification of proteomic heterogeneity

Considering the proteomic differences in spatially resolved regions within single tumors, we opt to quantitatively measure proteomics-based internal tumor heterogeneity (ITH). First, we constructed an internal variability score for each protein in each tumor using the proteomic data, which enabled the examination of the more variable and more stable processes. In agreement with the spatial tumor analysis, we found focal adhesion, keratins, and immune processes (phagocytosis and interferon-mediated signaling) to be among the most variable processes (Fig. 2A). In addition, metabolic processes, including cholesterol and carbohydrate metabolism presented high intratumor regional heterogeneity. High variability of immune processes might be associated with low immunotherapy response of breast cancer in general, as most tumors harbor regions with low immune signals. In the group of homogeneously-expressed proteins, we found enriched cell cycle-related, translation-related processes, as well as proteasome and ERAD pathways (Fig. 2B). These analyses suggest that similar to targeting clonal mutations, homogeneously-expressed proteins (e.g., cell cycle-related^15^ or proteasome-related^16^) may serve as potential therapeutic targets in combination with subtype-specific agents. We propose that these might be critical in achieving complete therapeutic responses. In contrast, targeting highly variable proteins may be insufficient to target the entire tumors.

**Fig. 2:**
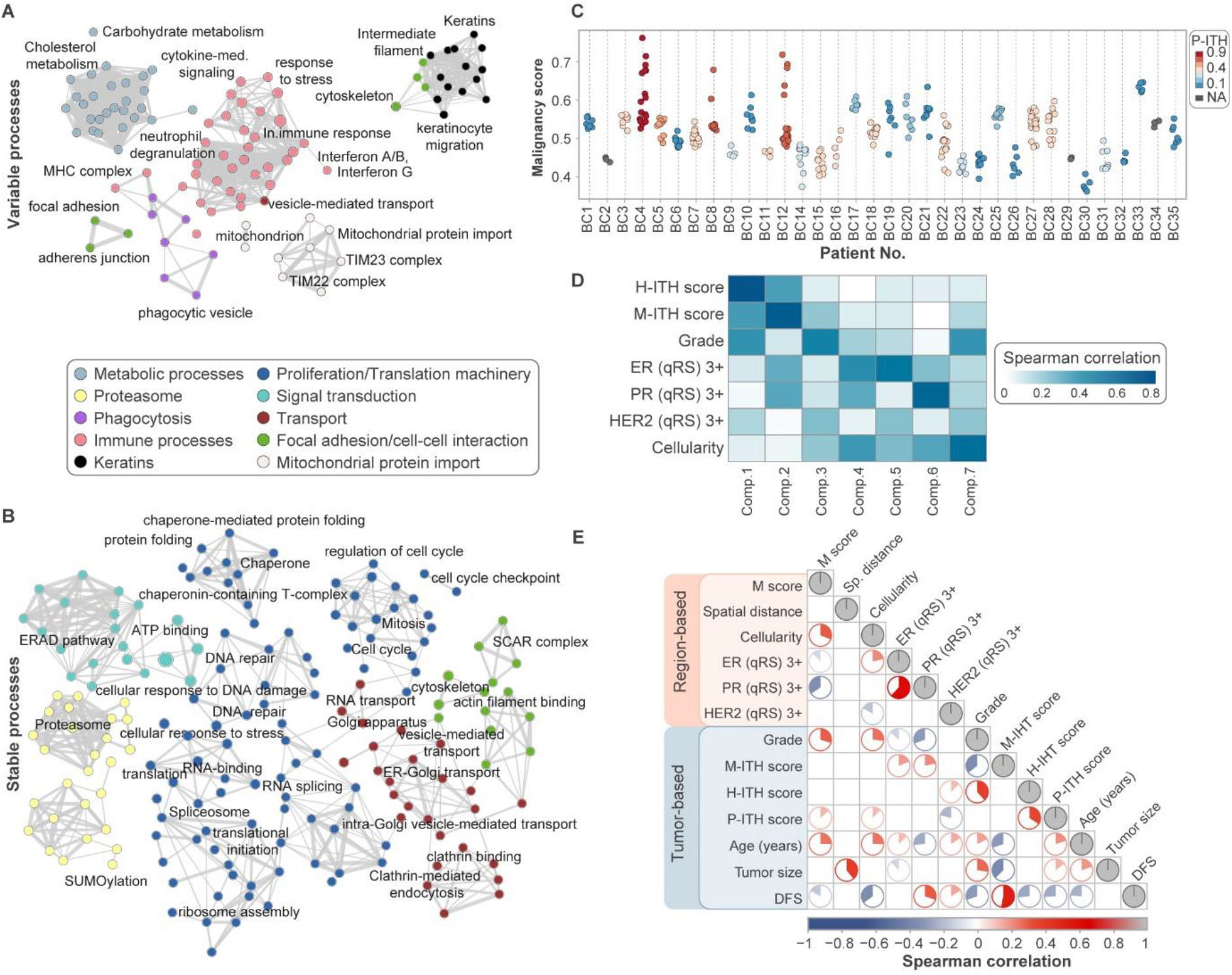
Proteomic-based intratumor heterogeneity. (**A-B**) Enrichment network analysis was performed separately for constant and variable proteins (based on the top 5% ranked proteins). It resulted in two networks with processes of variable proteins (top network) and constant proteins (bottom network) across all 35 tumors. Network nodes correspond to the enriched processes, edges represent mutual overlap. (**C**) Dotplot shows the intratumoral diversity of regional malignancy scores and interplay between two proteomics-based scores. Each tumor is color-coded based on its P-ITH score. (**D**) Correlation heatmap of seven canonical components to the pathological features shows that data variance is most predominantly explained by heterogeneity (H-ITH and M-ITH, first two components), then grade and molecular subtypes. Color scheme represents the results of absolute spearman rank correlation analysis. (**E**) Correlation heatmap of clinical (pathological) features and the calculated scores. Only the statistically significant correlations are shown (p < 0.05).

To complement the unsupervised variability analyses, we asked whether tumors possess homogeneous malignancy potential. Similar to the progression of disease approach, applied in genomic analyses^17^, we calculated a malignancy score per tumor region based on the correlation between its proteome and the average healthy control tissue (Extended Data Fig. 3A; see *Methods*). The malignancy score shows the extent of change from the healthy tissue and reflects the progression of a transformed phenotype, and was validated using grade of tumor progression (Extended Data Fig. 3B-D). Of note, while all regions within each tumor were defined as the same grade (excluding DCIS), most tumors showed a marked variance of malignancy scores of the different regions (Fig. 2C). Next, similar to genomic ITH scores, we calculated a proteomic heterogeneity score (P-ITH) per tumor as the average of two non-overlapping pairs of regions with the lowest correlation (Extended Data Fig. 3A; see *Methods*). To examine the interplay between the proteomics-based scores, we plotted the distribution of regional scores within each tumor and color-coded each tumor based on its P-ITH score (Fig. 2C). Tumors with highly variable proteomic features showed association with more diverging malignancy scores, which reflects the functional importance of the tumor proteomic heterogeneity.

### Molecular and histological subtype heterogeneity

Our histopathological tumor maps showed high variability of molecular subtypes (based on receptor expression) and histological subtypes. Sixty-six percent of the tumors in our cohort presented heterogeneous receptor expression levels that included HR+ (ER+/PR+/ERPR+) and TN regions (Extended Data Fig. 3E). Examination of the co-existence of distinct histological subtypes within single tumors showed that 17% of the tumors included more than one subtype (Extended Data Fig. 3F). Aiming to associate between the proteomic heterogeneity and the molecular and histological heterogeneity, we constructed additional heterogeneity scores based on pathological parameters: A molecular subtype heterogeneity score (M-ITH) and a histological subtype heterogeneity score (H-ITH). M-ITH and H-ITH scores were constructed similar to the Shannon diversity index^18^; however, we modified the formula to include the fraction of tumor area belonging to one subtype, and the number of coexisting subtypes within each tumor (Extended Data Fig. 3G; see *Methods*). Higher ITH scores correspond to a larger number of different subtypes within a single tumor and a lower dominance of a single subtype. We then examined the impact of the M- and H-ITH scores on the proteome. Canonical correlation analysis determined the degree of association between the proteomic data and the ITH scores and clinical features. The first two components, which explain the largest fraction of the variance, showed the highest correlation with H-ITH and M-ITH, respectively, and the third component highly correlated with tumor grade (Fig. 2D; Extended Data Fig. 4A). These results show the importance of the histopathologically heterogeneous nature of the tumors in shaping its proteome.

Next, we examined the interplay between all ITH scores and the malignancy score, defined above. Correlation analysis showed that malignancy scores correlate with P-ITH scores; tumor grade positively correlates with H-ITH (namely higher grades have higher heterogeneity of histological subtypes within single tumors), while having a negative correlation with M-ITH (Fig. 2E; Extended Data Fig. 4B). These findings suggest that histological subtypes may diverge with cancer progression, while molecular subtype heterogeneity reduces with progression, through intra-tumor competition or selection processes. Similarly, disease-free survival positively correlates with M-ITH, but negatively correlates with H-ITH and P-ITH; and the latter two highly positively correlated with each other (Fig. 2E). Overall, these results suggest distinct mechanisms associated with proteomic, histological, and molecular diversity during cancer progression.

### Functional landscapes of tumor heterogeneity

Given the significance of the internal tumor heterogeneity, we aimed to understand the functions associated with each of the heterogeneity scores. We extracted all proteins that correlated with each one of the four scores (Spearman rank correlation > 0.2) and constructed a joint protein-protein interaction network, in which each node is a protein, and the edges are the interaction confidence scores from the String database (Fig. 3A). Separate networks, colored for each score, reflected the higher levels of proliferation-related proteins and protein translation and turnover in the high malignancy scores. A mirror image, of lower proliferation-related proteins, was seen in the high M-ITH scores (in agreement with lower correlation with grade), with higher expression of metabolic and complement system proteins. Antigen presentation and cell cycle were enriched in high H-ITH; high IFNg and antigen presentation related proteins were enriched in high P-ITH scores (Extended Data Fig. 4C, D).

**Fig. 3:**
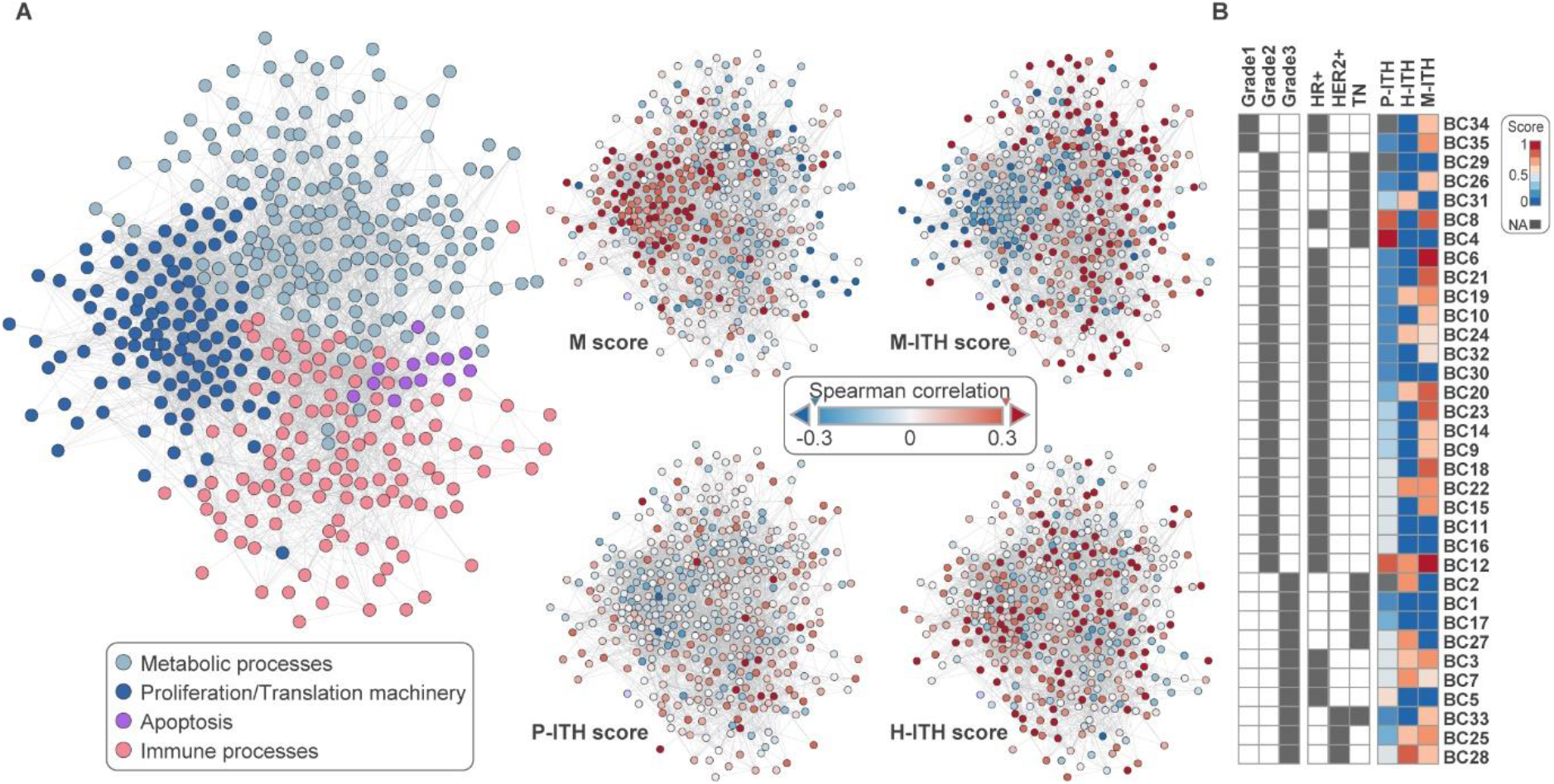
Analysis of multi-layered intratumor heterogeneity. (**A**) Merged protein-protein interaction network of the positively correlating proteins with each of the four scores (malignancy score and the three ITH scores, Pearson correlation > 0.2). On the left, the network is colored based on selected enriched annotations, and the four networks on the right are color-coded based on the correlation values with each one of the scores. (**B**) Integrated heatmap shows combination of clinical parameters and internal heterogeneity scores at the tumor level shows the switch between molecular and histological ITH depending on the tumor grade.

To account for subtype differences that may confound the previous analyses, we examined each molecular subtype separately (Fig. 3B). In accordance with the previous results, we found that homogeneous ERPR+, HER2+, and TN regions correlate with Grade 3, while when these molecular subtypes originated from heterogeneous tumors, they mostly correlated with Grade 2 (Extended Data Fig. 4E; see Methods). Therefore, the molecular heterogeneity is reduced with a higher grade, irrespective of the molecular subtype. In contrast, examination of the histological subtypes showed that homogeneous MPC, PLM, and ILC significantly correlated with Grade 2, while IDC-NST and MPC regions from heterogeneous tumors positively correlated with Grade 3 (Extended Data Fig. 4E). These results suggest that the histological subtypes (and specifically the combination of MPC and IDC-NST) diverge and are not selected against during breast cancer progression.

To associate between the heterogeneity status and all available clinical parameters and protein expression patterns, we performed unsupervised weighted gene co-expression network analysis (WGCNA; Extended Data Fig. 5). First, we observed a clear separation of the module correlation based on tumor grade (Fig. 4A; Extended Data Fig. 6A); modules with a strong positive correlation with Grade2 demonstrated a negative correlation with Grade 3, and vice-versa. In agreement with our previous results, the M-ITH score and Grade 2 mostly correlated with the same modules, which were enriched for mitochondrial metabolism, cell adhesion, and innate immunity (Fig. 4A). A closer look at the most highly correlating of these modules, MEturquoise, showed a high positive correlation with heterogeneous tumors of all subtypes, and negative correlations with high grade and TN-homogeneous tumors (Fig. 4B). In contrast to Grade2, in Grade 3 where we saw the increased molecular subtype homogeneity, none of the modules were common to both HR+ homogeneous and TN homogeneous regions, suggesting that distinct processes dominate tumor progression in a molecular subtype-specific manner.

**Fig. 4:**
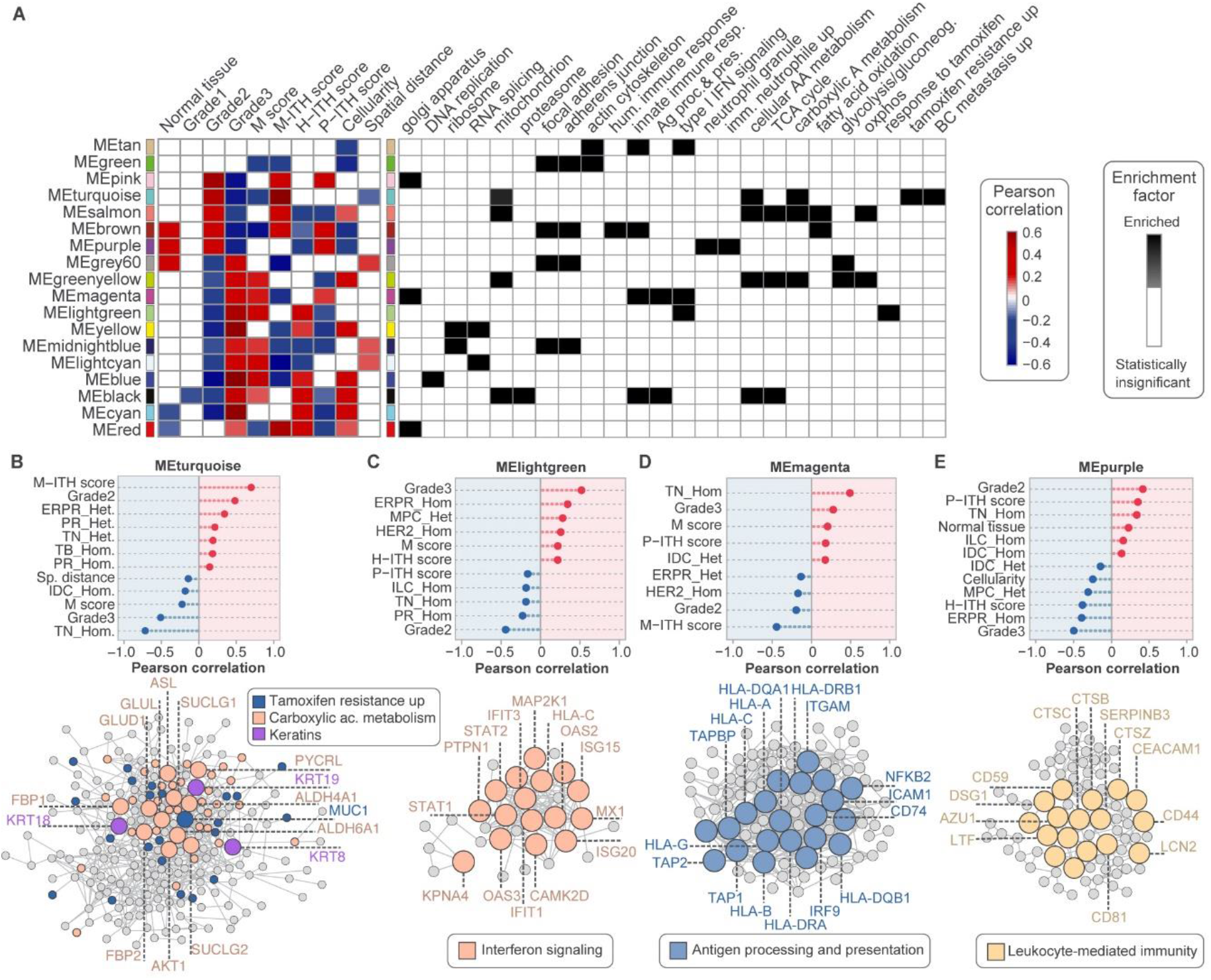
Unsupervised weighted gene co-expression network analysis (WGCNA) of different levels of intratumoral heterogeneity. (**A**) WGCNA of 321 regions (tumor and normal tissue) separated the proteomics data into 18 modules. The left heatmap shows the Pearson correlation of module eigengenes (MEs) with selected clinical parameters and heterogeneity scores. Only significant correlations are presented (Benjamini-Hochberg FDR q-value <0.05). The full correlation heatmap is presented in **Extended Data Fig. 6**. The right heatmap demonstrates functional enrichment for different MEs (Fisher exact test with Benjamini-Hochberg FDR q-value < 0.02). The statistical parameters of WGCNA are presented in **Extended Data Fig. 5**. **(B-E)** Cleveland plots show Pearson correlations of the pathological features to selected module eigengenes: MEturquoise (**B**), MElightgreen (**C**), MEmagenta (**D**) and MEpurple (**E**). Only significant correlations are presented (Benjamini-Hochberg FDR q-value < 0.05). Protein-protein interaction networks of the modules were generated based on the String database, and colored based on enriched KEGG pathways (Fisher exact test Benjamini-Hochberg FDR q-value < 0.02). See also **Supplementary Table 3C**.

### Impact of immune processes on tumor heterogeneity

Enrichment analysis of the WGCNA modules showed three modules, MElightgreen, MEmagenta, and MEpurple, enriched for immune processes. The MElightgreen module, which was enriched for IFN signaling, highly correlated with Grade 3, and homogeneous ERPR regions (Fig. 4C; Extended Data Fig. 6B), suggesting the involvement of anti-tumor immunity in eliciting high-grade homogeneity in this subtype. MEmagenta highly correlated with homogeneous TN tumors, heterogeneous IDC-NST, Grade 3 and high P-ITH, and negatively correlated with Grade 2 (Fig. 4D; Extended Data Fig. 6B). This module was enriched for antigen processing and presentation, and included multiple HLA proteins and *TAP* proteins, suggesting higher visibility to the immune system in high-grade TN tumors with coexisting proteomically-distinct regions and heterogeneous histological subtypes. MEpurple also positively correlated with homogeneous TN regions, however, it correlated with Grade 2 tumors and high proteomic heterogeneity, but negatively correlated with Grade 3, and ERPR homogeneous tumors (Fig. 4E; Extended Data Fig. 6B). Interestingly, this module was enriched for leukocyte-mediated immunity and included multiple immunosuppressive proteins, and checkpoint proteins, such as *CD81, CEACAM1*, and *CD59* (Fig. 4E; Extended Data Fig.6B). Thus, low grade, and specifically, Grade 2 homogeneous TN tumors may harbor immunosuppressive signals, that potentially retain high proteomic heterogeneity.

In agreement with the unsupervised WGCNA, a supervised comparison between heterogeneous and homogeneous tumors, while controlling for major confounding factors such as tumor grade and histological subtype showed that homogeneous grade 2 ERPR IDC-NST tumors present high immune-related proteins (*Stat, IFIT, HLA*) in the homogeneous ERPR+ tumors compared to heterogeneous ones, while regions from heterogeneous tumors presented higher levels of metabolic enzymes, related to glutathione metabolism, gluconeogenesis, and proline biosynthesis (Extended Data Fig.7A, C). A parallel analysis of TN regions showed higher proliferation-related proteins in the homogeneous tumors, including MCM complex proteins, while key TN markers (e.g., *KRT5, EGFR*) did not significantly differ according to the heterogeneity state (Extended Data Fig.7B-D). Of note, heterogeneous TN tumors also expressed markers of luminal subtypes (*KRT8, KRT18, MUC1*), despite being truly triple-negative according to immunohistochemistry (Extended Data Fig.7C-E). These results reinforce our findings that distinct pathways dominate homogeneous ERPR and TN tumors, while heterogeneous tumors of both ERPR+ and TN subtypes showed high proteomic similarity, with elevated expression of proteins related to glutathione metabolism and proline biosynthesis (Extended Data Fig.7B). Altogether, our results present the complex, multilayered heterogeneity and its impact on tumor progression and highlight the importance of broadening the current clinical classification in order to improve therapeutic potential.

## Discussion

Recent studies highlighted the necessity and urgency of a global tumor characterization approach in clinical practice by integrating different levels of information, especially for highly heterogeneous tumors^19–21^. Our proteomic analysis of histopathologically-defined regions created a multidimensional resource, which enables association between clinical features, spatial localization, and the proteomic landscapes in a systematic way. Using the supervised multi-region approach we demonstrated the advantage of capturing subtype-specific information and by adding spatial information of these regions, we found novel intratumor changes, among them, immune infiltrated tumor margins in TN regions, and elevated cell matrix adhesion in peripheral HR+ regions. These results might point at subtype-specific interactions within the tumor microenvironment^22–24^.

Our analyses identified high proteomic heterogeneity even in tumors that are otherwise defined as homogeneous based on histopathological parameters. Furthermore, the proteomic signals associated with tumor progression were found to markedly vary within single tumors, despite being defined as the same tumor grade. Interestingly, we found low correlation between the proteomic heterogeneity and the molecular subtype heterogeneity, suggesting that current clinical diagnostics may mask critical determinants of the tumor phenotype. We further found that reduced tumor heterogeneity is molecular subtype-dependent, dominated by cell proliferation and antigen processing and presentation, in triple-negative breast cancer tumors, and dominated by elevated interferon signaling in hormone receptor-positive tumors. These associations are only limited to the 35 patients in this cohort; however, they highlight potential critical regulations of tumor diversification and selection. In addition, our results revealed that TN regions from heterogeneous tumors have shared features with HR+ ones, suggesting that the microenvironment of these regions plays a more dominant role in shaping the proteome than the receptor expression level; these results testify to the risk of misdiagnosis when based on single needle biopsies^25,26^. In agreement, HR-negative cells, which express basal markers, within HR+ tumors were shown to confer resistance to standard endocrine and chemotherapies^27^.

Altogether, this study is the first to associate between the regional proteomic data and regulation of tumor heterogeneity, the involvement of the immune system, and intrinsic proliferative signals, in the context of the well-established breast cancer subtypes. In addition, our study can serve as a starting point for future basic and translational research. We envision that similar to targeting clonal mutations, targeting homogeneously-expressed proteins can be critical in reaching complete therapeutic responses. Furthermore, clinical assessment of tumor heterogeneity will be able to drive treatment decision-making, and will ultimately improve patient survival.

## Methods

### Clinical sample collection and tumor mapping

Formalin-fixed paraffin-embedded (FFPE) tissue samples were retrospectively collected from 37 breast cancer patients with primary tumors, with no pretreatment and various pathological characteristics. All samples were collected from the Institute of Pathology at the Sheba Medical Center and from the Israel National Biobank for Research (MIDGAM). The use of these samples for research was approved by the Institutional Review Board of Sheba Medical Center (approval SMC-7509-09) and the Tel Aviv University ethics committee, in accordance with the Declaration of Helsinki ethical guidelines. Tumor sections from five patients with hormone receptor-positive and triple negative breast cancer diagnosis were obtained from the MIDGAM according to protocol no. 130-2013, following approval by Ministry of Health. Clinical samples and clinical data were collected upon patient informed consent for collection, storage and distribution of samples and data for use in future research studies.

To map the histopathological characteristics of each tumor region, stained slides were assessed by trained breast pathologists from the Sheba Medical Center and receptor staining status was determined based on the Allred scoring system. To assess histological grade, H&E-stained slides were analyzed based on Elston and Ellis semi-quantitative method. Combining these scores, we created histopathological maps and defined distinct tumor regions according to the following criteria: receptor expression (per region), histological subtype (per region), and tumor grade (per region). These provide the visual representation of the intratumoral heterogeneity for downstream processing by laser capture microdissection (LCM). The tumor region cohort included invasive ductal carcinoma of no special type (IDC NST) of grade 1 (n=10), grade 2 (n=101) and grade3 (n=81), invasive lobular carcinoma (ILC) of grade 2 (n=29), pleomorphic carcinoma of grade 2 (n=27), micropapillary carcinoma of grade 2 (n=37), micropapillary carcinoma of grade 3 (n=22), tubular carcinoma of grade 2 (n=10), tubular carcinoma of grade 3 (n=4), papillary carcinoma of grade 3 (n=2), medullary carcinoma of grade 3 (n=7), non-invasive ductal/lobular carcinoma “in situ” (n=14) and normal glands (normal breast tissue) derived from tumor-free incision margin or contralateral mammary gland (n=19). Tumor regions defined based on receptor expression included: triple negative (TN) (n=98), hormone receptor-positive (HR+) (n=195), and HER2+ regions (n=18). All tumor-containing paraffin blocks were collected from every patient. The criteria for selection of regions for further processing was defined as 70% enrichment of cancer cells (within a region) with a requirement of minimal size of the target regions to provide sufficient material for proteomics (minimum 8 mm2). Detailed clinical information of each patient is included in Supplementary Table 1A-B. Disease-free survival was calculated from the date of primary treatment to the date of relapse or till the patient’s death.

### Immunohistochemistry (IHC) staining

Five consecutive 3.5 μm tissue sections were sliced from all cancer-containing paraffin blocks from each patient (average 4-5 blocks per patient), mounted on positively charged glass slides, and dried overnight at 37°C. Slides were mapped by staining with H&E, anti-ER (DAKO), anti-PR (DAKO), anti-HER2 (DAKO), and anti-cytokeratin (Cell Marque). Staining was performed on the BOND-RX automated staining platform using Bond Polymer Refine Detection automated kit (Leica Biosystems). H&E staining was performed manually using a standard protocol.

### Image analysis

Stained slides were scanned using the Leica Aperio VERSA Digital Pathology Scanner (Aperio Technologies Inc.). Staining quantification of all selected regions was performed using the eSlide Manager software via the Aperio image analysis algorithms: Nuclear (ER, PR), Membrane (HER2), and Cytoplasmic (CK). Regions were defined by a pathologist and manually annotated in the software. Regions were considered to be positive for the antibody staining if they reached a cutoff threshold of 10% and defined as high-intensity staining (+3). Immune cell infiltration per region was quantified by modified nuclear algorithm via the eSlide Manager software.

### Laser capture microdissection (LCM)

PEN-membrane glass slides were used to allow combined UV and IR laser microdissection. Eight μm sections were mounted on the slides, dried and deparaffinized with xylene, followed by a series of graded ethanol. Staining was performed with the Paradise Plus system (Thermo Fisher Scientific). To obtain sufficient protein amounts for downstream analyses, we selected a minimum tissue area of 8 mm2 for each region of interest (corresponding to approximately 13,000 cells and 2 μg protein). Stained tumor samples were microdissected using the ArcturusXT laser capture microdissection system (Thermo Fisher Scientific). LCM system allowed us to harvest relatively pure target populations of cancer cells using precise ultraviolet (UV) and infrared (IR) lasers under direct microscopic visualization.

### Proteomics sample preparation

Microdissected tumor sections were collected in Eppendorf LoBind microcentrifuge tubes, lysed, and digested using two different protocols, using trifluoroethanol (TFE) or SDS in two sample batches. Protocol I included lysis with 50% 2,2,2-TFE in 50 mM ammonium bicarbonate buffer, the samples were boiled at 95°C for 1 h, sonicated for 10 cycles in a Bioruptor sonicator (Diagenode), and then centrifuged at 17000 xG for 20 min to pellet tissue debris. Lysates were transferred to fresh LoBind tubes, reduced with 5 mM dithiothreitol (DTT), and alkylated with 15 mM iodoacetamide (IAA) for 30 minutes, followed by overnight “in-solution” digestion using LysC-trypsin mix as described^1^. Protocol II included lysis in 4% Sodium dodecyl sulfate (SDS) in 25 mM HEPES (pH=8) lysis buffer, incubation for 1.5 h at 95°C, reduction, alkylation, and digestion following the Single Pot Solid Phase Sample Preparation (SP3) protocol^2^. SeraMag Hydrophilic and Hydrophobic beads (GE Healthcare) mix was used at a concentration of 100 μg/μl for overnight “on-bead” digestion using LysC-trypsin mix (Promega)^2,3^. Tryptic peptides from 363 regions were labeled using 10plex TMT reagent and assembled into 41 TMT-labeled sample sets. Each set was composed of nine channels for the different regions and one channel for an internal standard. The internal standard was always labeled with the 131-C reagent. Assembled TMT-labeled peptides were then fractionated offline into eight fractions using high pH reverse-phase chromatography (PierceTM High pH Reversed-Phase Peptide Fractionation kit), following the manufacturer instructions. Fractionation was followed by LC-MS analysis.

### LC-MS-based proteomics

LC-MS/MS analysis was performed using high-performance liquid chromatography (Easy nLC 1000 HPLC system) coupled on-line to a Q-Exactive HF mass spectrometer (Thermo Fisher Scientific) through the EASY-Spray ionization source. Peptides were separated on 75 μm × 50 cm long EASY-spray PepMap columns (Thermo Fisher Scientific) and loaded with Buffer A (0.1% formic acid). Peptides were eluted with a 140-minute gradient of water-acetonitrile, and each batch of samples, composed of eight factions, was analyzed in a total of 20 hours. We slightly modified the gradients for each fraction as follows: for fractions 1 and 2, 5-30% buffer B (80% acetonitrile/0.1% formic acid); for fraction 3, 5-31% buffer B; for fraction 4, 5-32% buffer B; for fraction 5, 5-34% buffer B; for fraction 6, 5-36% Buffer B; for fraction 7-8, 5-38% buffer B. All fractions were eluted at a flow rate of 200 nl/min at 40°C. MS data were acquired in a data-dependent mode and the acquisition method included a full scan event at a resolution of 120,000, followed by top 10 MS/MS scans at a resolution of 60,000.

### Data Processing: proteomics raw MS data processing

Raw MS files were analyzed with the MaxQuant software (version 1.6.2.6) and the Andromeda search engine^4^. The Uniprot database (human, version from 2018 with 95057 entries) was used for searching MS/MS spectra. Modifications for 10plex TMT labels were defined in MaxQuant and included the label impurities as indicated for each label batch. A decoy database was used to determine a 1% false discovery rate for both protein identification and peptide spectrum matches. We included carbamidomethyl cysteine as a fixed modification and N-terminal acetylation and methionine oxidation as variable modifications with a maximum of five modifications per peptide. Maximum two missed cleavages were allowed.

### Proteomic statistical analysis

All statistical tests were performed using Perseus, Python, and R. We used the proteinGroups output table for the analysis (Supplementary Table 2). Initial data processing included filtration of proteins originating from the decoy database (reverse proteins), proteins only identified by modification sites and common contaminants. In addition, we filtered out extracellular matrix proteins to avoid distortion of the overall ratio distribution by high and variable expression of extracellular proteins. The 33 regions that had less than 3000 protein IDs were excluded from the analysis. Protein expression values (reporter intensities corrected) were log2-transformed and normalized to the internal control in each TMT set. Then the entire dataset was filtered to retain proteins with a minimum of 70% of valid values, reaching 4046 protein groups, which were used for most downstream analyses. The data were further zero-centered by subtracting the median of each sample. We imputed missing values by creating an artificial normal distribution from the missing values, with a downshift of 1.9 standard deviations and a width of 0.5 of the original ratio distributions, and after that, applied quantile normalization algorithm using R code from the “Biobase” package. The entire cohort was analyzed in four batches, including two sample preparation procedures, as described above. To correct the batch effect of the two sample preparation procedures, we subtracted the first component in a Principal Component Analysis. For supervised analyses, all normalizations were performed on the selected group of samples included in each specific analysis. This allowed maximizing the number of proteins included in each analysis. Normalizations included Log2-transformation, normalization to the internal control, filtration to retain proteins with at least 70% valid values, and zero-centering.

Specific statistical analyses were performed as follows:

- Analysis of pseudo-bulk samples was performed by calculating the median expression of each protein from regions of the same tumor. Subtype annotations for the bulk tumors were based on the clinical reports of the whole tumor. For comparison of keratin expression levels between pseudo-bulk and region data, we performed Kruskal–Wallis test and created boxplots using “ggpubr” package in R. Significantly differentially expressed proteins between pseudo-bulk subtypes were extracted by performing Welch’s test with Benjamini-Hochberg (BH) FDR of 0.1. In order to compare the results with multi-sampled regions and correct for the significant differences in sample size, Welch’s test (BH FDR 0.1) was performed by drawing random samples (the same number of samples as in the pseudo-bulk analysis) from the dataset. The random sampling was repeated 100 times and the number of significantly different proteins was annotated.
- **Unsupervised clustering** was performed using modified minimum variance method (‘ward.D2’). For the analysis, we used non-imputed matrix with 100% valid values and calculated Euclidean distances between regions. Then applied the modified Ward’s method to create groups with minimized variance within clusters. To represent our cohort of 330 regions, we used Tree-and-Leaf dendrogram where each node represents a tumor region, branches of the tree constructed based on the Euclidean distances and cluster separation presented by spatial localization of the branches. For visualization we used R code from the “TreeAndLeaf” package.
- **Supervised analysis** based on subtype and grade was performed using one-way ANOVA followed by post Hoc tests with permutation-based FDR cutoff of 0.05 using the Perseus platform. Results were visualized in R using the ‘ComplexHeatmap’ package. Enrichment analyses were performed using Fisher exact test with an FDR cutoff of 0.05, and the entire dataset of 8150 proteins as background. To demonstrate variation of malignancy scores we used the “ggplot2” package in R for visualization.
- **Canonical correlation analysis (CCA)** was performed to reduce dimensionality of the proteomics data. First, we used scikit-learn PCA in Python to reduce the data dimension to 240 components, which explained >99% of the variance. Second, we performed canonical correlation analysis of the proteomics and clinical data, further reducing the data to its 7 canonical correlation components. For the analysis, we used 309 tumor regions (excluded normal tissue) which have receptor staining quantification data. For visualization of the CCA results, we plotted 3D scatterplots using “scatterplot3d” package in R. To show the data hierarchy, we used absolute Spearman rank correlation analysis between canonical components and pathological features, and visualized it using R code from the “pheatmap” package.
- **Spatial distance calculation** was performed by measuring the physical distance from the tumor margin to each region of the same tumor. To obtain measurements, we used the eSlide Manager software via the Aperio image analysis tools (ruler). First, we calculated the absolute spatial distance (absSpaD), and then normalized each distance to the tumor size to construct normalized spatial distance (nSpaD) to compare across tumors.

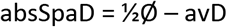

where ½Ø corresponds to the half of the measured tumor diameter and avD is the average distance from the tumor margin Normalized spatial distance (nSpaD) is the distance from the tumor center, normalized to the tumor size:

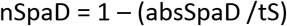

where tS corresponds to the tumor size.
- **Stable and variable proteins calculation** was calculated as the median absolute deviation (MAD) scores per protein. Based on the MAD score we ranked every protein per tumor. Using sum of ranked proteins, we selected top 5% for stable and variable group and performed functional enrichment. We visualized the output in Cytoscape 3.8.2 via the “EnrichmentMap” plugin. For the EnrichmentMap, we used node FDR cutoff of 0.05 and edge cutoff (Jaccard similarity index) of 0.25. Redundant processes were removed from resulted networks.
- **Score calculation** of intratumor heterogeneity on different levels generated three internal tumor heterogeneity (ITH) scores and a malignancy score. For molecular and histological subtypes, we calculated the number of coexisting subtypes per tumor, and then obtained the area measure of each subtype via the eSlide Manager software. The score was calculated as

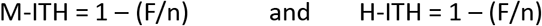

where F corresponds to the largest fraction of subtype from the total area of tumor and n is number of subtypes coexisting in the same tumor. Obtained scores were calculated per tumor. For the proteomic ITH score, we calculated the correlations between all pairs of tumor regions using log2 transformed non-normalized dataset of 8150 protein groups, and then averaged the data from two non-overlapping pairs with the lowest correlation per tumor to avoid any subjective effect of multisampling. The score was calculated per tumor which allowed us to avoid batch effect and opt for using a whole non-normalized dataset of 8150 proteins. The score was calculated as

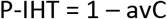

where avC corresponds to average proteome correlation within a tumor. Obtained scores were calculated per tumor. For malignancy score calculation we applied analysis of disease progression by averaging the proteomic data of normal regions of 13 patients from our cohort, and then calculated the Pearson correlation of each tumor region to the normal tissue. Log2 transformed normalized data of 4046 protein groups was used as input. The scores were calculated as

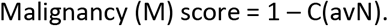

where C(avN) corresponds to proteomic correlation to averaged normal regions. Obtained scores were calculated per region.
- **Conversion of heterogeneity status (based on M-ITH and H-ITH scores) from the tumor to the region level** was done for each region and defined whether it is homogeneous or heterogeneous (molecularly or histologically), in the context of the entire tumor. Each region was assessed based on its subtype definition and whether its own subtype definition was different/same as the majority of the tumor area (>70% of the area).
- **To create protein networks correlating to the scores**, first, we filtered proteins based on their Pearson correlation to each score with established cutoff of 0.2. After merging the lists, we built a protein network using STRING output via Cytoscape. 1D annotation enrichment analysis was performed on the score values with a Benjamini–Hochberg FDR threshold of 0.02 using Perseus software. A Chi-square test was performed for comparative correlation between tumor grade and heterogeneity of the tumors using R code from the “stats” package.
- **Weighted gene co-expression network analysis (WGCNA)** was performed using R code from the “WGCNA” package using signed network, bicor correlation function, power = 10 and a reassign threshold of 0.1. Out of 330 regions, six regions were excluded as outliers and three regions were excluded due to the absence of receptor quantification data. Results of the WGCNA were visualized using “pheatmap” and “WGCNA” packages in R. For enrichment analysis we performed Fisher exact test with a Benjamini–Hochberg FDR q-value < 0.02 using Perseus software. To validate our findings in a supervised manner, while controlling for major confounding factors such as tumor grade and histological subtype, we selected only Grade 2 tumors of one histological subtype (IDC-NST). A Welch’s test with permutation-based FDR cutoff of 0.05, comparing between homogeneous and heterogeneous status of each subtype was performed using Perseus software on non-imputed data.
- **Clinical information and staining quantification** results of the assembled cohort were visualized using R code via the ‘ComplexHeatmap’ package. Receptor staining quantification and clinical data correlations were visualized using “pheatmap”, “corrgram” and “corrplot” packages in R.

### Data and software availability

All raw data are available via ProteomeXchange with identifier PXD024190. Reviewer account details: *Username:*; *Password:* xUr5MpUo

## Acknowledgements

The Geiger laboratory received Funding from the European Research Council (ERC) starting grant [639534] and the Israel Science Foundation (grant 748/16). We thank Georgina Barnabas, Gali Yanovich-Arad, Michal Harel, Michael Selitrennik, and the members of Geiger laboratory for the fruitful discussions and assessment of the manuscript. We thank Eilam Yeini and Professor Ronit Sachi-Fainaro for help with immunohistochemistry; Dr. Adi Zundelevich for assistance with patient cohort assembly; members of the Pathology Department at the Sheba Medical Center for assistance in tissue selection and processing; Israel National Biobank for Research (MIDGAM) for assistance in tissue selection. We thank Constantiner Institute for the partial support.

## Author contributions

Conceptualization, T.G, and M.M; Investigation, M.M; Formal analysis, M.M, A.S, R.S, D.P, N.I and I.B; Data curation, K.K, D.N, I.M and B.S; Resources, I.B and E.N.G; Writing Original Draft, M.M and T.G; Writing Review & Editing, M.M, A.S, R.S, E.N.G and T.G; Supervision, T.G; Funding acquisition, T.G.

## Materials & Correspondence

Tamar Geiger

## EXTENDED DATA FIGURES

**Extended Data Fig. 1:**
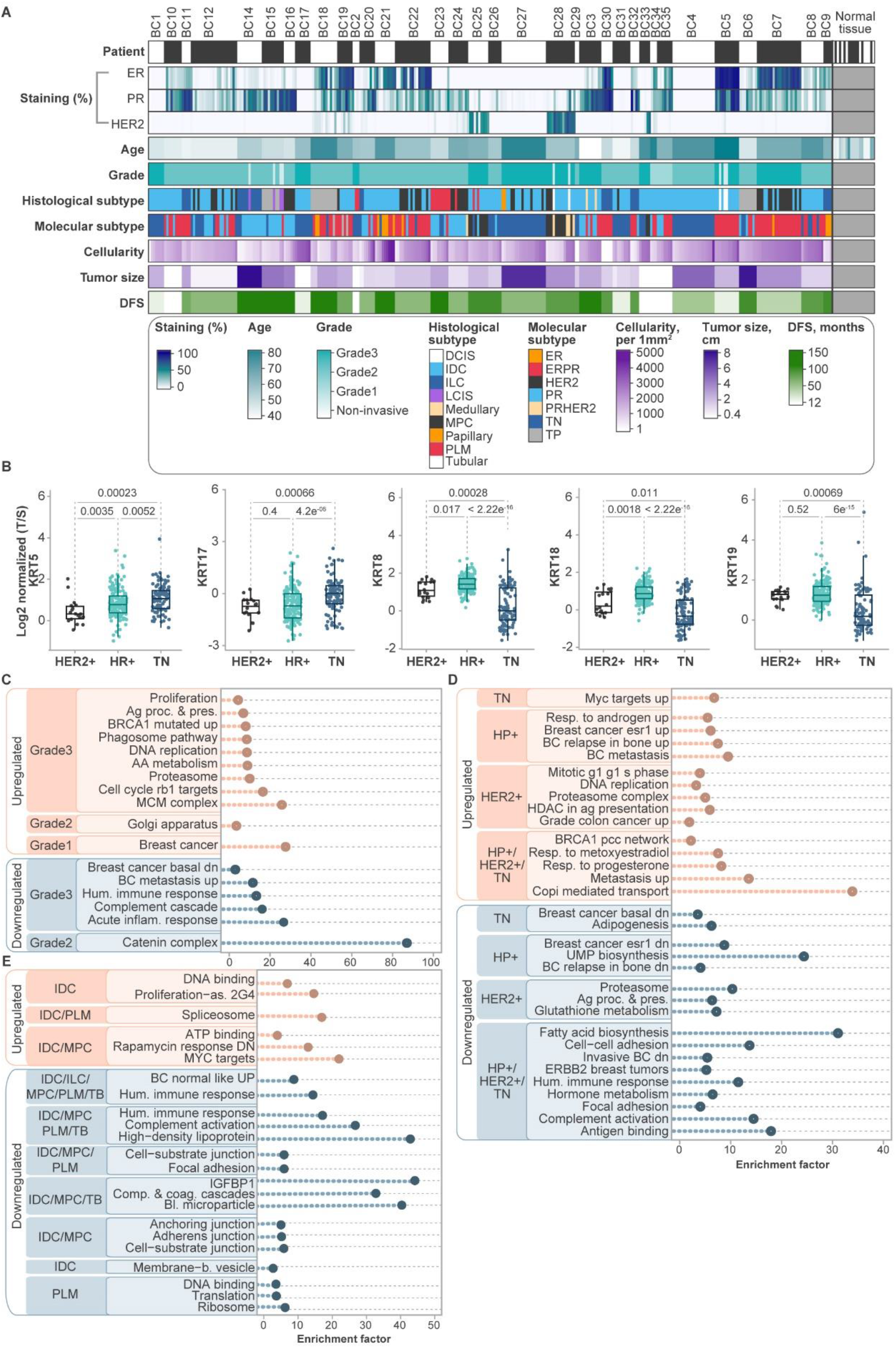
Clinical features of the cohort and proteomic comparison to healthy control samples. (**A**) The study cohort includes 330 regions derived from 35 breast cancer patients. Heatmap indicates the clinical parameters. Patient row separates between all patients by switching black and white bars, and all normal samples in the cohort are indicated in grey. (**B**) Boxplots demonstrate subtype specific changes of keratin expression in the different regions. Statistical significance was determined with Kruskal–Wallis test, p<0.05. HR+ regions include ER, ERPR and PR+ regions. (**C**) Enrichment analyses of selected processes that deviate from healthy tissue show processes associated with tumor grade. The entire list of enriched processes can be found in the **Supplementary Table 3A**. (**D**) Enrichment analyses of selected processes that deviate from healthy tissue show processes associated with molecular subtype. The entire list of enriched processes can be found in the **Supplementary Table 3A**. (**E**) Enrichment analyses of selected processes that deviate from healthy tissue show processes associated with histological subtype. The entire list of enriched processes can be found in the **Supplementary Table 3A**.

**Extended Data Fig. 2:**
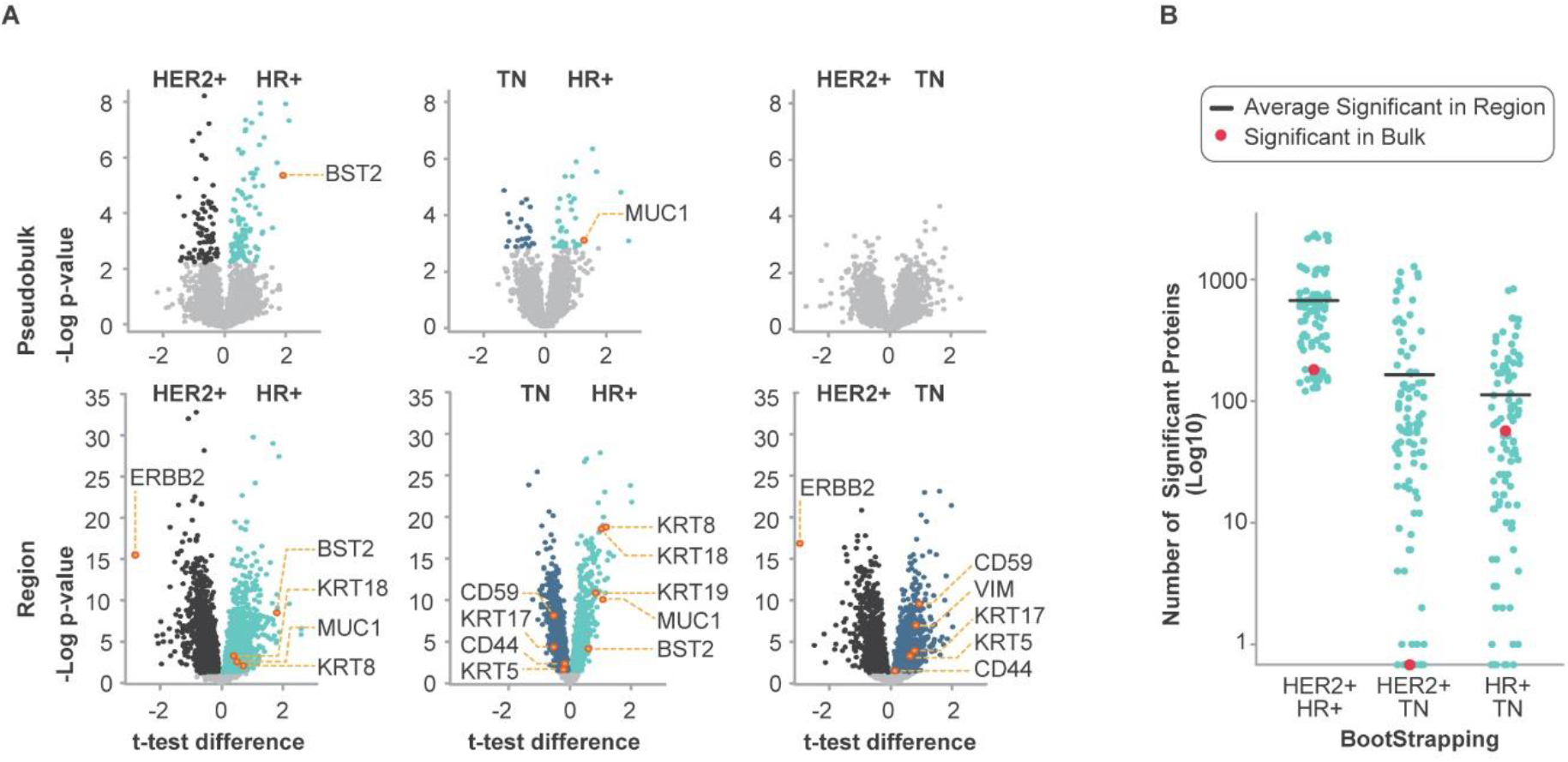
Advantage of multi-region approach for breast cancer analysis. (**A**) Volcano plots show the advantage of regional data over bulk analyses in capturing subtype specific significantly changing proteins. Statistical significance was determined with Welch’s t-tests, with permutation-based FDR q-value < 0.05. (**B**) Dotplot presents the number of significantly different proteins upon bootstrapped Welch’s test (with 100 iterations of random sampling). The number of significantly changing proteins in the pseudo-bulk analysis is indicated by a red dot.

**Extended Data Fig. 3:**
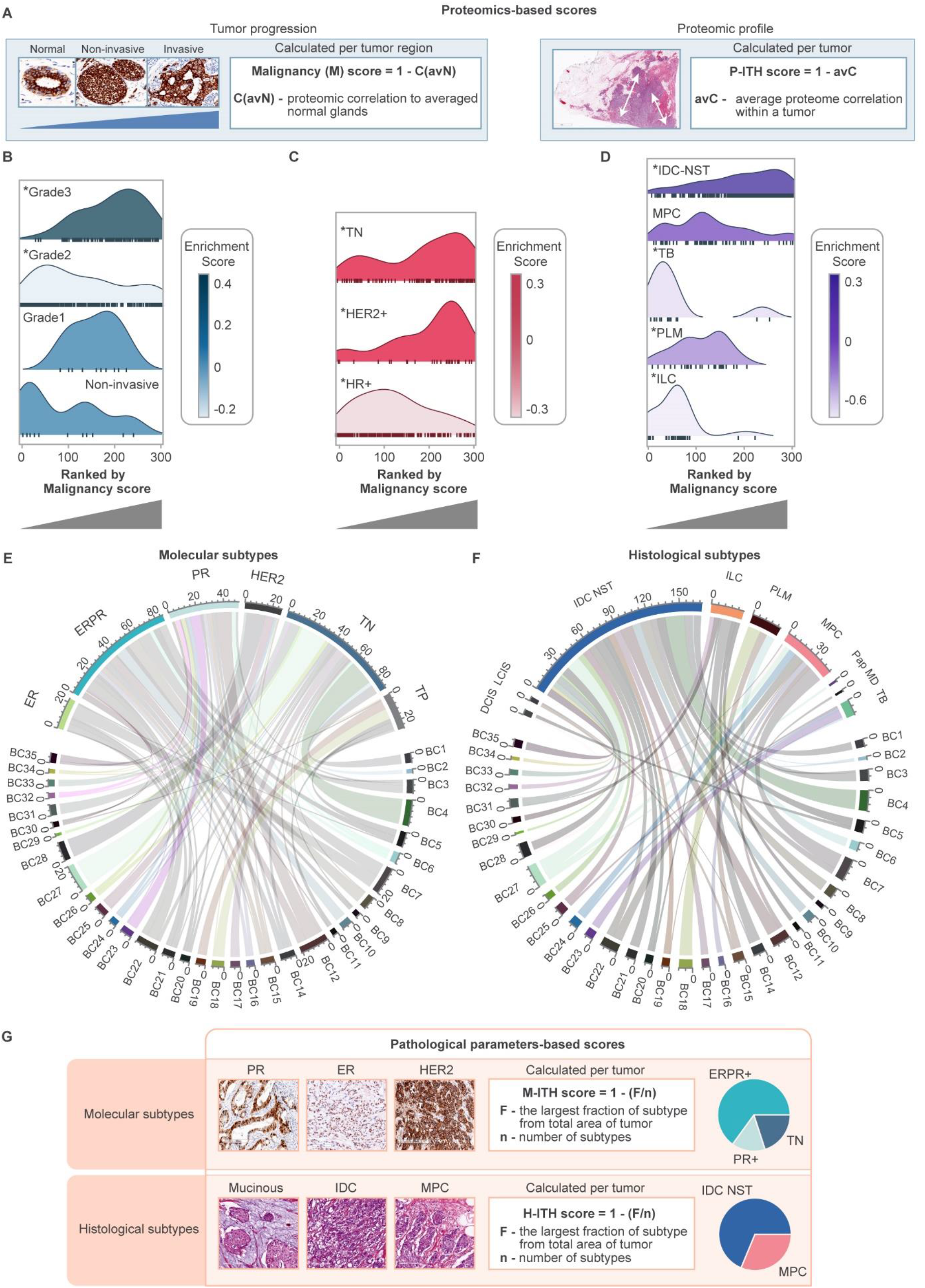
Analysis of intratumor heterogeneity. (**A**) Quantitative analysis of proteomics-based intratumoral heterogeneity defined malignancy score (M) and proteomic heterogeneity score (P-ITH) based on the proteomic data. M score is calculated per region, while P-ITH score is calculated per tumor. (**B-D**) Ridgeline density plots demonstrate ranking of tumor regions by their malignancy scores. Higher malignancy scores are presented in Grade 3 tumor regions (**B**), in TN and HER2 regions (**C**), and in IDC-NST regions (**D**). Grade 2 tumors present a wide distribution of malignancy scores. Asterisks represents FDR q-value < 0.05. (**E, F**) Chordplots show distribution of regions of each tumor between represented subtypes: Molecular (**E**) and histological (**F**). Each tumor consists of several regions that belong to one or several subtypes. HR+ regions (ER, ERPR, PR) demonstrate higher intratumoral heterogeneity than TN regions. Plots show that most tumors are composed of more than one histological and more than one molecular subtype. (**G**) Quantitative analysis of pathological intratumoral heterogeneity defined molecular subtypes heterogeneity score (M-ITH) and histological subtypes heterogeneity score (H-ITH). M-ITH and H-ITH are based on clinical parameters (H&E and receptor IHC); both scores are calculated per tumor.

**Extended Data Fig. 4:**
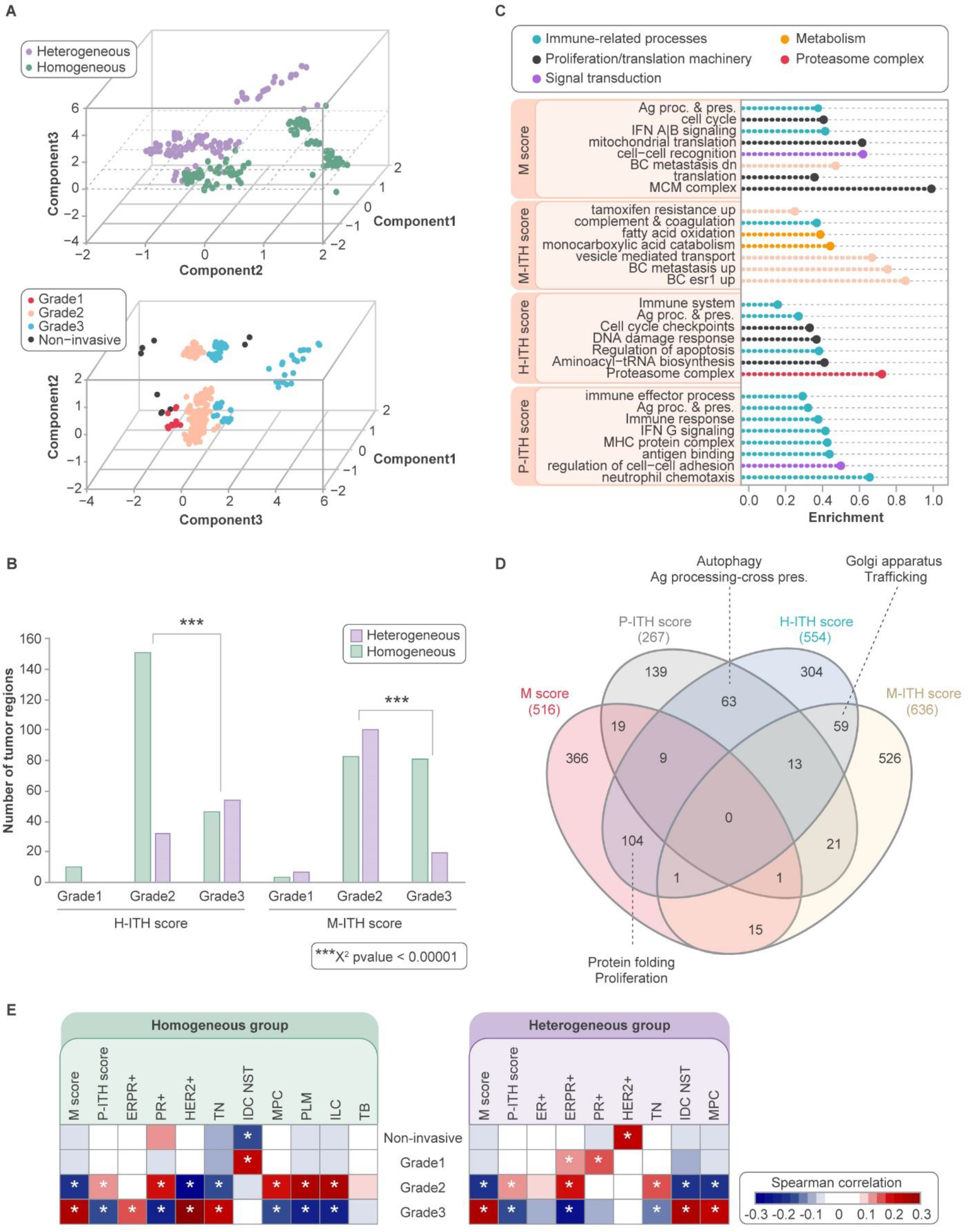
Analysis of data hierarchy and heterogeneity scores. (**A**) 3D scatter plots of the first three components of the canonical correlation analysis of the 309 tumor regions data were segregated based on their histological and molecular heterogeneity (top scatterplot) and tumor grade (bottom scatterplot). (**B**) Barplot shows the separation of the different tumor grades in the groups of histological and molecular homogeneous and heterogeneous tumors ***p < 0.00001. (**C**) Cleveland plot shows the functional enrichments of four scores (Benjamini-Hochberg FDR q-value < 0.02). The entire list of enriched processes can be found in the **Supplementary Table 3B**. (**D**) A Venn diagram shows the overlap of positively correlating proteins with each one of the scores. (**E**) Correlation heatmap of clinical parameters separated based on the heterogeneity status of molecular and histological subtypes. On the left heatmap, homogeneous tumors and on the right heatmap heterogeneous tumors. Asterisk represents p < 0.05.

**Extended Data Fig. 5:**
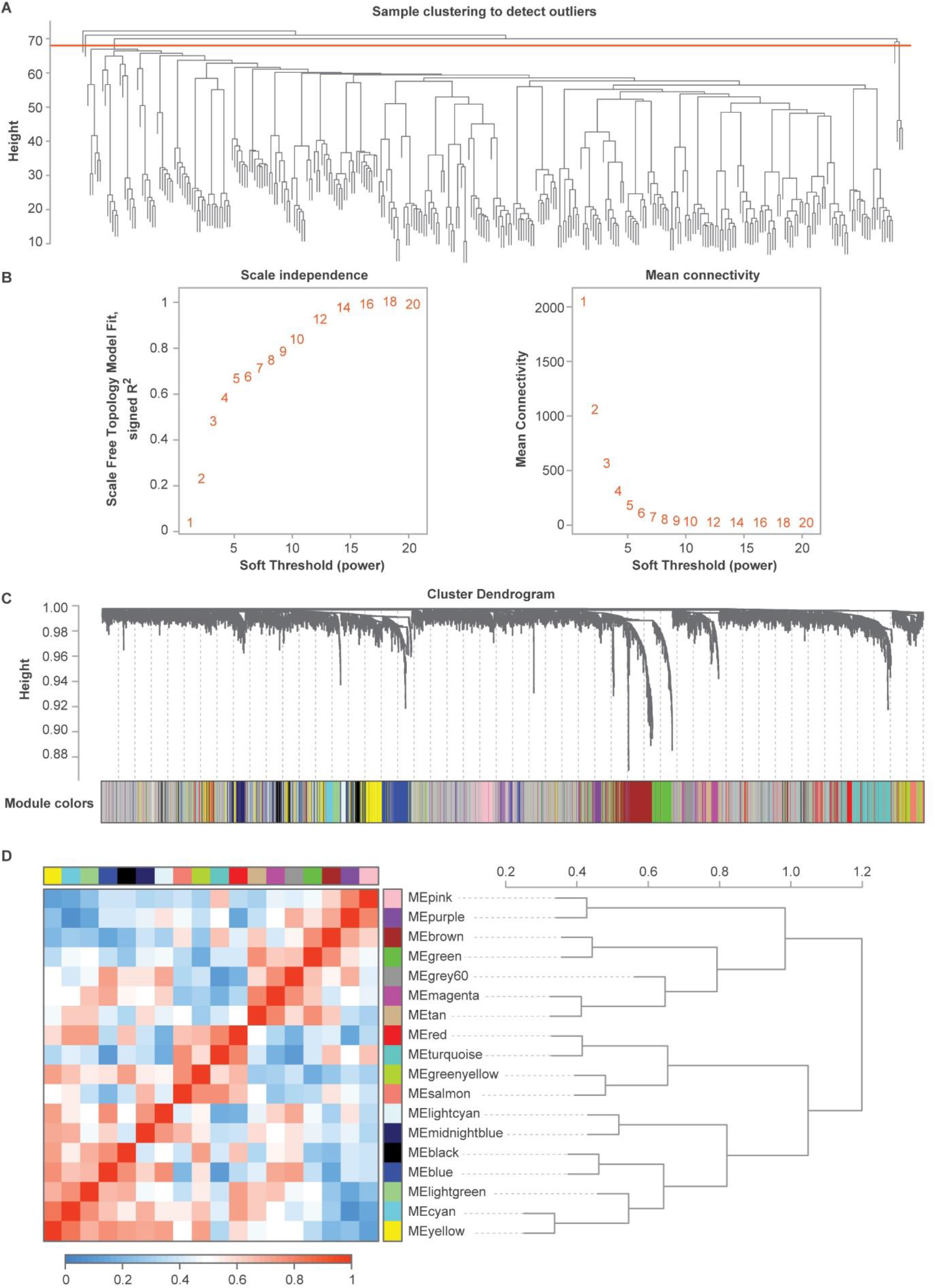
WGCNA analysis. (**A**) Clustering of 330 regions within the WGCNA to remove outliers (red line). Six outlier samples were removed. Three additional samples were removed due to missing annotations. (**B**) Summary network indices (y-axes) as functions of the soft thresholding power (x-axes), where the numbers in the plots indicate the corresponding soft thresholding powers. The plots indicate that approximate scale-free topology is attained around the soft-thresholding power of 10. (**C**) Protein dendrogram obtained by clustering the dissimilarity based on consensus Topological Overlap with the corresponding module colors indicated by the color row. Consensus eigengene networks and its differential analysis shown by clustering tree of the consensus module eigengenes and heatmap with high adjacency and low adjacency to each module.

**Extended Data Fig. 6:**
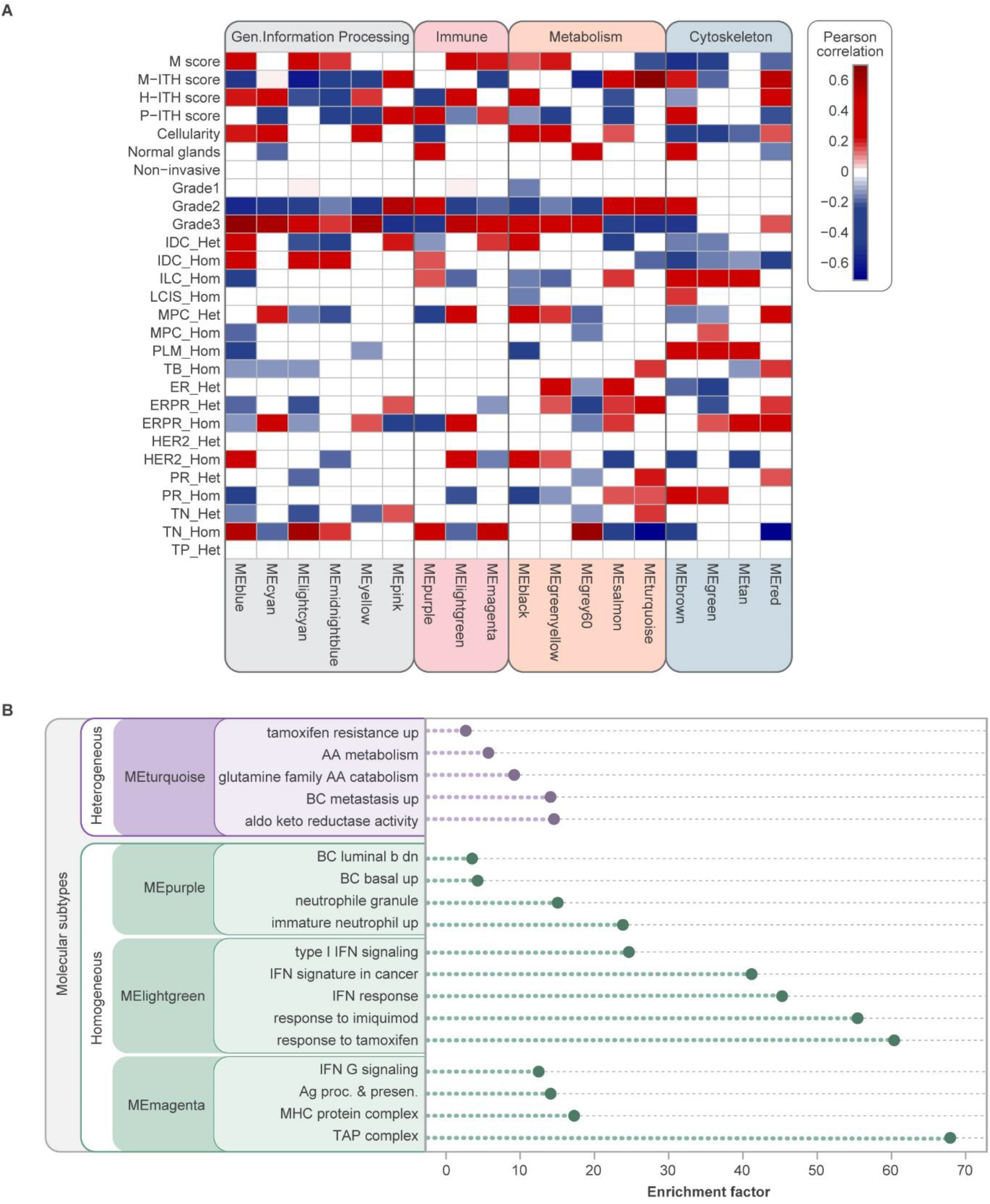
WGCNA of region proteomic data. (**A**) WGCNA of 321 regions (tumor and normal tissue) shows module eigengenes (MEs) significantly correlating with different levels of heterogeneity and tumor grade (extended heatmap, related to **Fig. 4**). Only significant correlations are indicated (Pearson correlation, FDR q-value < 0.05). (**B**) Cleveland plot shows the functional enrichment of selected modules (FDR q-value < 0.02). The entire list of enriched processes is given in the **Supplementary Table 3C**.

**Extended Data Fig. 7:**
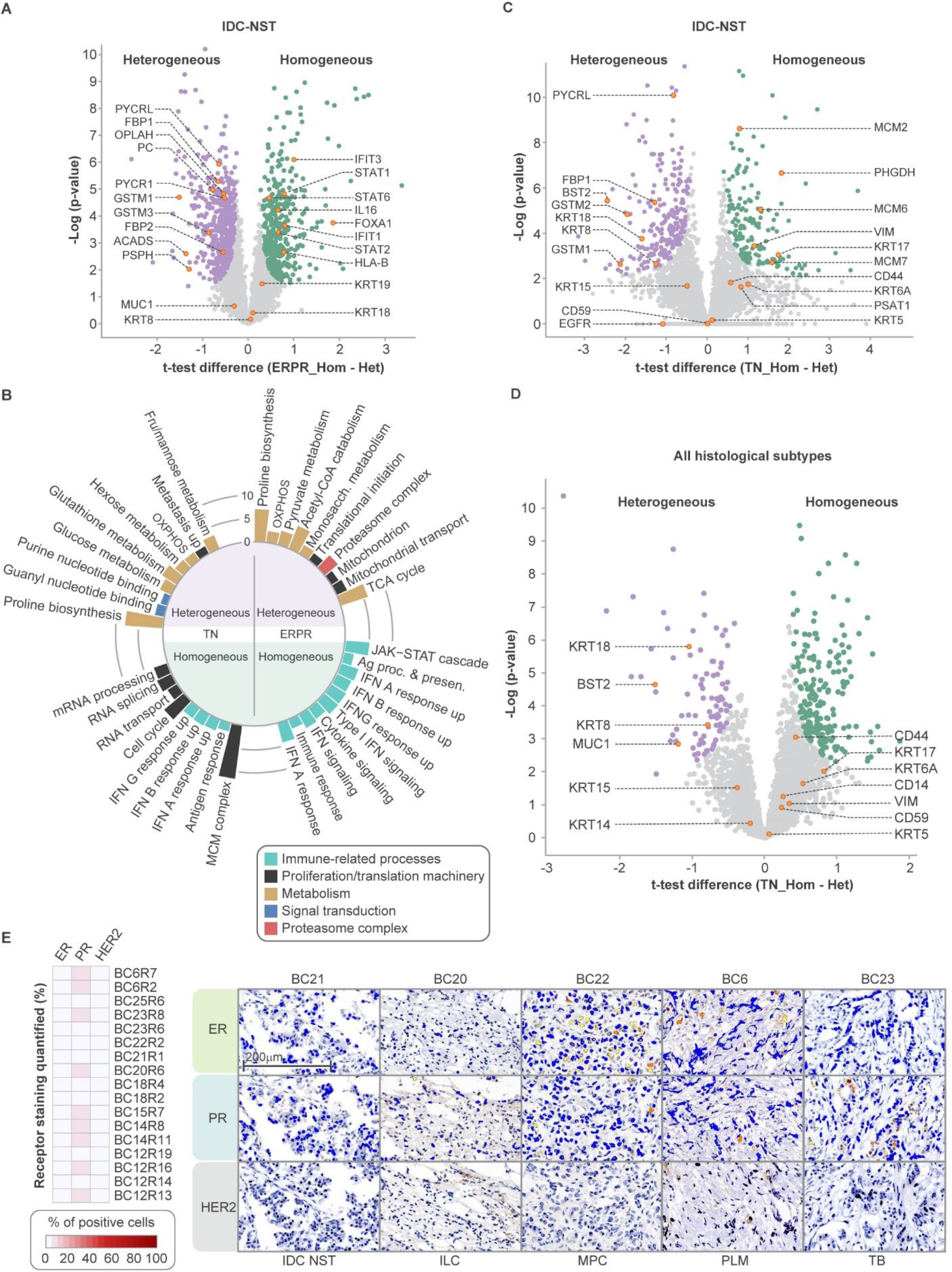
Functional differences between heterogeneous and homogeneous regions. (**A**) Volcano plots demonstrate the results of Welch’s test comparing homogeneous and heterogeneous groups based on molecular heterogeneity for ERPR+. The comparison was performed between 20 regions derived from five patients (heterogeneous group) and 12 regions derived from two patients (homogeneous group). (**B**) Enrichment analysis of significantly changing proteins from (**A**) and (**B**) (Fisher exact test Benjamini-Hochberg FDR q-value < 0.02). Circular bar plot presents the enrichment factor. The entire list of enriched processes can be found in **Supplementary Table 3D**. (**C**) Volcano plot shows the results of Welch’s test comparing homogeneous and heterogeneous TN regions. Markers of basal subtype indicated below the cutoff threshold (grey); biomarkers of luminal subtype indicated in the heterogeneous group (above the cutoff threshold). The comparison was performed between five regions from two patients (heterogeneous group) and 36 regions from five patients (homogeneous group). (**D**) Volcano plot shows the results of Welch’s test comparing homogeneous and heterogeneous TN regions. The comparison is similar to the one shown in (**C**), except the regions of grade 2 are represented by all the histological subtypes across the cohort. The comparison was performed between 17 regions from nine patients (heterogeneous group) and 37 regions from five patients (homogeneous group). (**E**) Percentages of cells stained positive for ER, PR and HER2 and their combinations in the panel of selected 17 TN regions from heterogeneous group that were used in (**D**). Examples of receptor staining per histological subtypes from heterogeneous group of TN regions.

